# Individual variation in phenotypic plasticity of the stress axis

**DOI:** 10.1101/600825

**Authors:** Sarah Guindre-Parker, Andrew G. Mcadam, Freya Van Kesteren, Rupert Palme, Rudy Boonstra, Stan Boutin, Jeffrey E. Lane, Ben Dantzer

**Affiliations:** Department of Integrative Biology, University of Guelph, Ontario, CANADA; Department of Psychology, University of Michigan, Michigan, USA; Department of Biomedical Sciences, University of Veterinary Medicine, Vienna, AUSTRIA; Department of Biological Sciences, University of Toronto Scarborough, Ontario, CANADA; Department of Biological Sciences, University of Alberta, Alberta, CANADA; Department of Biology, University of Saskatchewan, Saskatchewan, CANADA; Department of Ecology and Evolutionary Biology, University of Michigan, Michigan, USA

**Keywords:** Faecal glucocorticoid metabolites, density, adaptive plasticity, red squirrels

## Abstract

Phenotypic plasticity—one individual’s capacity for phenotypic variation under different environments—is critical for organisms facing fluctuating conditions within their lifetime. North American red squirrels (*Tamiasciurus hudsonicus*) experience drastic among-year fluctuations in conspecific density. This shapes juvenile competition over vacant territories and overwinter survival. To help young cope with competition at high densities, mothers can increase offspring growth rates via a glucocorticoid-mediated maternal effect. However, this effect is only adaptive under high densities, and faster growth often comes at a cost to longevity. While experiments have demonstrated that red squirrels can adjust hormones in response to fluctuating density, the degree to which mothers differ in their ability to regulate glucocorticoids across changing densities remains unknown—little is known about within-individual plasticity in endocrine traits relative to among-individual variation. Findings from our reaction norm approach revealed significant individual variation not only in a female red squirrel’s mean endocrine phenotype, but also in endocrine plasticity in response to changes in local density. Future work on the proximate and ultimate drivers of variation in the plasticity of endocrine traits and maternal effects is needed, particularly in free-living animals experiencing fluctuating environments.

## INTRODUCTION

All organisms experience changes in their environment, and the ability to adjust morphology, physiology or behaviour according to environmental conditions can provide individuals with important fitness benefits [1]. Phenotypic plasticity—when one individual or genotype can produce multiple phenotypes across a gradient of environments—is thought to represent an important mechanism that allows organisms to respond to environmental changes [2]. Phenotypic plasticity may be particularly important for organisms in fine-grained environments [1], defined by spatial or temporal fluctuations of key environmental features that occur within an individual’s lifespan [3,4].

North American red squirrels (*Tamiasciurus hudsonicus*, hereafter, ‘red squirrels’) experience drastic fluctuations in their fine-grained environment, where an important aspect of a red squirrel’s environment—local conspecific density—can vary up to 4-fold within an individual’s lifetime [5]. Periods of high density pose a challenge for breeding individuals, where the availability of vacant territories critical for offspring overwinter survival is low and competition for these vacancies is high [6–8]. Red squirrel mothers can prepare their young to cope with high density conditions via an adaptive hormone-mediated maternal effect [5]: under high densities, mothers with elevated glucocorticoids during pregnancy give birth to faster growing pups that have a greater probability of surviving their first winter [5]. This maternal effect is only adaptive under high densities [5,9], however, since faster growth does not improve juvenile recruitment under low density [5,10]. Offspring born under high densities have shorter lifespans [10], suggesting an inverse link between growth rate and lifespan [5]—this would be consistent with fitness costs of compensatory growth in mammals [11–13]. Chronically elevated glucocorticoids could also have negative impacts on mothers, leading to oxidative stress [14], immunosuppression [15,16], and reduced parental care [17]. As a result, we would expect the optimal red squirrel maternal phenotype to include elevated glucocorticoids during periods of high density, but decreased glucocorticoids during periods of low density. While glucocorticoids are positively related to naturally and experimentally elevated density in red squirrels [5], the degree to which individual mothers vary in their endocrine plasticity in response to changes in density remains unclear [18–20].

Glucocorticoids are broad mediators of phenotypic plasticity in vertebrates [19,20], circulating at flexible concentrations [21] and promoting physiological and behavioural adjustments following perturbations in an animal’s environment [18,22–24]. This endocrine trait itself is plastic [25,26], where individuals can regulate concentrations of glucocorticoids in response to diverse environmental stressors [27,28]. Here, we take a reaction norm approach to explore within-individual variation in glucocorticoid plasticity in red squirrel mothers experiencing drastic among-year environmental fluctuations. We measured faecal cortisol metabolites (FCM)—a non-invasive measure of adrenocortical activity [29]—to determine whether female red squirrels show (i) individual variation in FCM and/or (ii) individual variation in plasticity of FCM in response to density changes.

## METHODS

### Field data collection

We studied two unmanipulated populations (‘Kloo’ and ‘Sulphur’) that have been monitored since 1987, as part of the Kluane Red Squirrel Project in the southwestern Yukon in Canada (61° N, 138° W). Each individual red squirrel defends an exclusive territory containing a hoard of white spruce (*Picea glauca*) cones (called a ‘midden’) over their lifespan. Seeds from cached cones sustain squirrels through winter, making territory-ownership crucial for survival [6–8]. Individuals were uniquely marked with numbered ear tags threaded with a unique combination of coloured wires. Territory ownership was assessed reliably each spring via a population-wide census, described in detail elsewhere [30]. Briefly, populations were completely enumerated each year and territory ownership was confirmed via a combination of observations of territorial vocalizations and live-trapping (by placing food-baited Tomahawk Live Traps on middens, Tomahawk, WI, USA). We calculated each individual’s local population density in the spring as the number of neighbours owning a midden within the acoustic environment of the focal individual (i.e. within a 130 meter radius) [31].

Between 2006 and 2014, we collected faecal samples opportunistically when trapping individuals (from February to September, mean time of day at capture ± S.D. = 11:30 a.m. ± 3 hours). We checked below traps for fresh faeces, which we kept on ice until they could be frozen (within five hours of collection) [32]. At trapping, we assessed each female’s breeding status (pregnant, lactating, or non-breeding) by palpating the abdomen for foetuses and checking nipple condition [33]. Faecal cortisol metabolites were assayed in one of two facilities (Michigan and Toronto), following identical protocols which have been validated previously [32,34]. A subset of samples (*N* = 128) analysed in both labs showed a strong positive correlation (Pearson correlation = 0.88), suggesting variation among labs had minimal effects on our results. Samples were thawed, lyophilized, flash-frozen, and pulverized by mortar and pestle. Steroids were extracted using methanol (1 mL of 80% methanol for 0.05 grams of dry faeces) [32,35], and the supernatant was used in an enzyme immunoassay to quantify glucocorticoid metabolites with a 5α-3β,11β-diol structure [32]. Using a sample quality control run on all assay plates (*N* = 115), we found that estimates of optical density for these were highly repeatable (*R* = 0.85, 95% confidence intervals = 0.54 - 0.93). Faecal cortisol metabolites are expressed as ng/g of dry faeces, and ln-transformed to meet the assumptions of parametric statistical tests.

### Statistical analyses

Our dataset included 1,729 FCM measurements, collected from 153 females, where 57 individuals had repeated FCM measurements across a range of densities (i.e. two or more years). We did not censor individuals sampled in only one year, which reduces statistical power [36]— including females sampled in a single year in our models helps to parameterize fixed effects and does not bias estimates for variation in plasticity (i.e. random slopes).

To determine if female red squirrels differed in their endocrine plasticity we used a random regression modelling approach [36,37], fitting four general linear mixed models (LMMs) by maximum likelihood, and identifying the best supported model(s) using Akaike’s information criterion [38,39]. We performed LMMs in R [40] using the package ‘lme4’ (version 1.1-17). Diagnostic plots revealed that model residuals were normally distributed and were not heteroscedastic. We identified model(s) most strongly supported given our dataset using the ‘bbmle’ package (version 1.0.20) to calculate model AIC_C_ scores and model weights. Lower AIC_C_ scores indicate stronger support, and models within two AIC_C_ values of one another fit a dataset similarly well [39].

We first built a null model including the following variables known to influence FCM in red squirrels [5,32]: breeding status, linear and quadratic effects of Julian date, the lab where samples were analysed (i.e. assay ID), and a linear effect of local spring density as fixed effects. There did not appear to be any nonlinear effects of any predictor beyond the quadratic effect of sampling date. We standardized continuous fixed effects (i.e. with mean = 0, S.D. = 1), and checked for multicollinearity (all variance inflation factors were below 2). The null model did not include random effects. The second model we tested was a random intercept model which added a random intercept for each individual squirrel to the null model. The last two models we tested added a random slope for density to the random intercept model, which allowed us to test for individual variation in endocrine plasticity as local density changed across years [36]. The third model assumed there was no correlation between random intercepts and slopes, whereas the fourth model allowed for random intercepts and slopes to be correlated. This tested whether an individual’s endocrine phenotype affected the likelihood that it would exhibit weaker or stronger plasticity.

## RESULTS

We identified two equivalent top models that had a combined weight of 99%, and both included random intercepts and slopes (models 3 and 4; Table 1). Model 4 also included a negative correlation between individual estimates for intercepts and slopes, though this correlation was not statistically significant (the 95% confidence intervals overlapped with 0; Table 2). The other two models had a ΔAIC of 8 or more, indicating they were not supported by the data (Table 1).

**Table 1:** We compared four candidate LMMs fit with maximum likelihood differing in their random effect structure in order to test for individual differences in endocrine plasticity. Fixed effects were identical in all models (i.e. breeding status, assay ID, linear & quadratic Julian date of sample collection, and density).

| Model | Random Effects | Covariance<br>(intercepts & slopes) | *df* | *AIC<sub>c</sub>* | $\Delta AIC$ | Model Weight |
| --- | --- | --- | --- | --- | --- | --- |
| Model 1: Null | n/a | n/a | 8 | 3670.4 | 142.3 | <0.001 |
| Model 2: With ID | Intercept (ID) | n/a | 9 | 3536.3 | 8.1 | 0.009 |
| Model 3: With ID x density, no covariance | Intercept (ID)<br>Slopes (ID x density) | No | 10 | 3528.4 | 0.2 | 0.469 |
| Model 4: With ID x density, with covariance | Intercept (ID)<br>Slopes (ID x density) | Yes | 11 | 3528.2 | 0.0 | 0.522 |

**Table 2:** We identified two top models, for which we present the estimates and *t*-values of fixed effects, variances and standard deviations of random effects, and the correlation coefficient for the covariance between individual intercepts and slopes (model 4 only). We estimated 95% confidence intervals (CI) by parametric bootstrap using the ‘lme4’ package. While we do not report *P*-values, parameters with 95% CI not overlapping with 0 can be considered as significantly different from 0 (indicated with an asterisk)—this is consistent with *P*-values calculated using the Satterthwaite approximation in ‘lmerTest’ (version 3.1-0).

| Fixed Effects | Model 3<br>(ID x density, no covariance) |  |  | Model 4<br>(ID x density, with covariance) |  |  |
| --- | --- | --- | --- | --- | --- | --- |
|  | Estimate ± SE | *t*-value | 95% CI (Estimate) | Estimate ± SE | *t*-value | 95% CI (Estimate) |
| Intercept (lactating, Toronto) | 7.90 ± 0.05 | 161.3 | 7.08 – 8.00 * | 7.90 ± 0.05 | 159.8 | 7.80 – 8.00 * |
| Assay ID (Michigan) | -0.01 ± 0.04 | -0.394 | -0.09 – 0.06 | -0.02 ± 0.04 | -0.405 | -0.10 – 0.07 |
| Breeding Status (non-breeding) | 0.12 ± 0.05 | 2.327 | 0.02 – 0.20 * | 0.10 ± 0.05 | 2.273 | 0.01 – 0.20 * |
| Breeding Status (pregnant) | 0.09 ± 0.05 | 1.880 | 0.002 – 0.18 * | 0.09 ± 0.05 | 1.909 | -0.007 – 0.19 |
| Julian Date | -0.04 ± 0.02 | -2.489 | -0.08 – -0.01 * | -0.04 ± 0.02 | -2.404 | -0.07 – -0.004 * |
| Julian Date <sup>2</sup> | -0.06 ± 0.02 | -3.937 | -0.09 – -0.03 * | -0.06 ± 0.02 | -3.926 | -0.10 – -0.03 * |
| Density | 0.15 ± 0.03 | 5.011 | 0.09 – 0.21 * | 0.13 ± 0.03 | 4.297 | 0.07 – 0.19 * |
| Random Effects | Variance | Standard Deviation | 95% CI (Standard Deviation) | Variance | Standard Deviation | 95% CI (Standard Deviation) |
| ID (intercept) | 0.079 | 0.281 | 0.22 – 0.33 * | 0.079 | 0.281 | 0.21 – 0.35 * |
| ID x Density (slope) | 0.028 | 0.168 | 0.07 – 0.22 * | 0.038 | 0.195 | 0.11 – 0.26 * |
| Residual | 0.397 | 0.630 | 0.60 – 0.65 * | 0.395 | 0.629 | 0.61 – 0.65 * |
| Covariance of Random Effects | Correlation Coefficient | 95% CI (Correlation) |  | Correlation Coefficient | 95% CI (Correlation) |  |
| Intercept (ID) & Slope (ID x Density) | NA | NA |  | -0.37 | -0.74 – 0.02 |  |

The top models support that female red squirrels show individual variation in FCM, as well as individual variation in endocrine plasticity across changes in density (Figure 1). We did not find support for a correlation between random intercepts and slopes (Table 2). Thus, an individual’s tendency to have elevated FCM was independent of the degree to which they exhibited endocrine plasticity. The fixed effects in both models supported previous findings in this system, where FCM increased with local density and declined non-linearly with Julian date (Table 2) [5,32].

**Figure 1:**
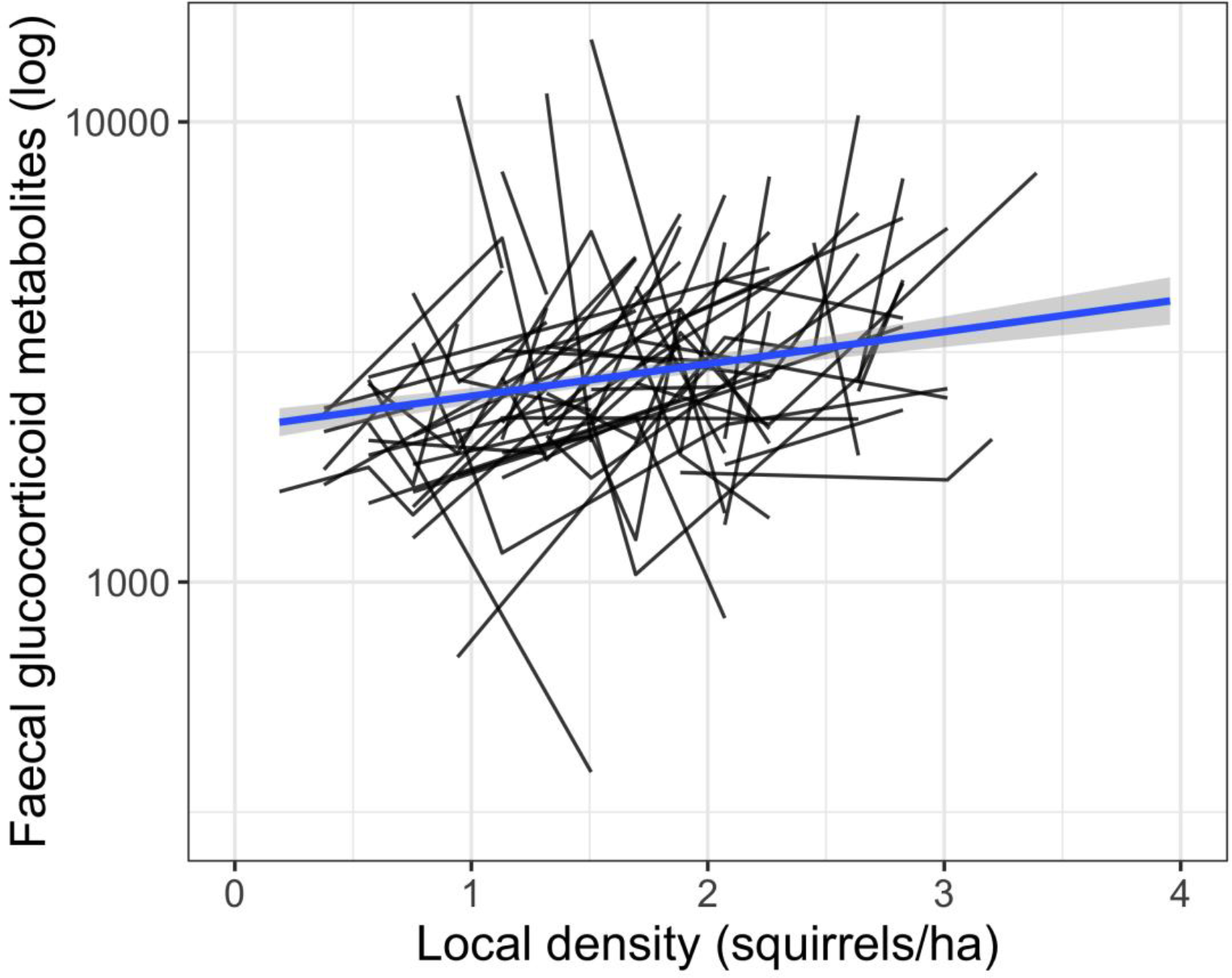
Reaction norms for all repeatedly-sampled female North American red squirrels (*N* = 57). Each black line indicates one individual’s reaction norm, connecting mean annual endocrine phenotype across years (i.e. changes in population density). The blue line indicates the population-wide relationship between faecal glucocorticoid metabolite concentrations and local density.

## DISCUSSION

This study highlights three key results about endocrine variation in a free-living population of red squirrels. First, individuals differed consistently in FCM. Second, females differed in their endocrine plasticity in response to changes in local population density. Over half of females had elevated FCM as population density increased (56% of females), whereas 9% of females showed little change and 35% of females showed a decline in FCM with increasing density. Finally, our results suggest that an individual’s mean FCM phenotype does not covary with their plasticity in FCM in response to changes in their social environment.

Our results add to a growing body of literature that show significant individual variation in glucocorticoid plasticity. House sparrows (*Passer domesticus*) showed individual variation in the degree to which glucocorticoids declined with age [41], or with food availability [25]. In both cases, some individuals responded strongly to changes in age or food availability, whereas others showed little endocrine response. Similarly, free-living male chimpanzees (*Pan troglodytes*) showed repeatable individual variation in urinary glucocorticoid responses to circadian changes [42]. Our study is the first to examine variation in endocrine plasticity along a natural gradient of ecological conditions, which provides important insight into how organisms differ in their ability to track changes in their environments. Future studies characterizing endocrine plasticity in free-living animals will be critical to better predict how individuals, populations, or species may cope with changing environments.

The prevalence of individual variation in glucocorticoid plasticity across studies suggests that individuals frequently differ in their abilities to respond to changes in environmental conditions, though the proximate mechanism underlying these differences remains unknown. In red squirrels, individuals that do not increase glucocorticoids under elevated densities could have responded plastically in downstream targets of glucocorticoids (e.g. changing receptor densities or corticosteroid-binding globulins [43]). A second possibility is that individuals showing little change in glucocorticoids across densities may simply be constrained in their ability to regulate glucocorticoid secretion [18]. For example, animals with elevated glucocorticoids may already be operating at their physiological maximum and may not be able to increase circulating concentrations further; if this were the case, however, we would expect to find a negative correlation between intercepts and slopes (which was not supported). A third possibility is that individuals differ in their ability to perceive local density—individuals underestimating density could fail to upregulate glucocorticoids under high-density conditions.

More broadly, it is unclear whether individual differences in endocrine plasticity arise from genetic, early-life, or environmental effects. Circulating glucocorticoids are shaped in part by additive genetic effects [44–47], though the heritability of plasticity in glucocorticoids has not been examined [18]. Early life exposure to fluctuating environments [48], maternal glucocorticoids [49], and reduced parental care [50] all shape the glucocorticoid phenotype of offspring [51–53], and could similarly shape variation in endocrine plasticity. Future research on endocrine plasticity is needed to understand (i) the proximate mechanism that generates variation in glucocorticoid plasticity, and (ii) the evolutionary causes and consequences of variation in glucocorticoid plasticity.

## Acknowledgments

We thank the Champagne and Aishihik First Nations for allowing us to conduct research within their traditional territory. We are also grateful to the teams who contributed to data collection and analysis. This is publication number XX of the Kluane Red Squirrel Project.

## Author Contributions

BJD, AGM and SGP conceived of the research question. BJD, SB, AGM, and JEL supervised and funded the field data collection. BJD, FVK, RP, and RB contributed to the endocrine lab work. SGP conducted analyses with guidance from AGM and BJD. SGP, AGM, and BJD wrote the first draft of the manuscript and all authors contributed to subsequent drafts.

## Data Accessibility

We will deposit data in Dryad, following a 5-year embargo.

## Ethical Statement

This research was approved by the University of Michigan (PRO00005866, PRO00007805), the University of Guelph (AUP09R006, AUP1807) and Michigan State University (AUF 03/05-033-00).

